# Identification of RNA-Binding Protein Targets with HyperTRIBE in *Saccharomyces cerevisiae*

**DOI:** 10.1101/2023.03.17.532881

**Authors:** Weilan Piao, Chong Li, Pengkun Sun, Miaomiao Yang, Yansong Ding, Wei Song, Yunxiao Jia, Liqun Yu, Hua Jin

## Abstract

As a mater regulator in cells, RNA-binding protein (RBP) plays critical roles in organismal development, metabolism and various diseases. It controls gene expression at multiple levels mostly by specific recognition of target RNA. The traditional CLIP-seq method to detect transcriptome-wide RNA targets of RBP is less efficient in yeasts due to their cell walls. Here, we established an efficient HyperTRIBE (Targets of RNA-binding proteins Identified By Editing) in yeast, by fusing a RBP to the hyper active catalytic domain of human RNA editing enzyme ADAR2 and expressing the fusion protein in yeast cells. The target transcripts of RBP were marked with new RNA editing events and identified by high-throughput sequencing. We successfully applied TRIBE to identifying the RNA targets of two yeast RBPs, KHD1 and BFR1. The antibody-free HyperTRIBE has competitive advantages including low background, high sensitivity and reproducibility, and a simple library preparation procedure, which provides a reliable strategy for RBP target identification in *Saccharomyces cerevisiae*.

## Introduction

RNA-binding protein (RBP) specifically interacts with RNA in cells, involved widely in the regulation of gene expression^1^. RBP-RNA interaction makes the basis of cell functions, thus, the investigation of RBP-RNA interaction network is of great significance for comprehending different cellular processes and understanding the fundamental roles of RBP in these processes. Hundreds of RBPs have been estimated to occur in mammals^2,3^, and multiple RBPs usually co-work as a complex and some proteins in the complex can be associated with RNA indirectly, which gives rise to the difficulties in identification of RNA-protein interaction^4,5^.

Given the importance of RNA-RBP interaction, the methods profiling the target RNAs of a given RBP at a transcriptomic level have been developed rapidly and the representative methods include RNA-immunoprecipitaion sequencing (RIP-seq), high-throughput sequencing of RNA isolated by crosslinking immunoprecipitation (HITS-CLIP) and its variants, TRIBE, HyperTRIBE, RNA tagging, and Surveying Targets by APOBEC-Mediated Profiling (STAMP)^6–12^. HITS-CLIP (CLIP-seq) is a powerful method for detecting RBP-binding sites on RNAs at a single-nucleotide resolution and has been widely used^9^. Although original HITS-CLIP relied on high-quality antibodies and a large number of cells with low UV-crosslinking efficiency, its variants have greatly improved the efficiency^10–12^.

In recent years, three antibody-independent methods have been developed by fusing a RNA modification enzyme to a RBP respectively to mark the RBP targets, providing alternative strategies for the studies of RBPs. TRIBE and RNA tagging developed in late 2015 to early 2016 and STAMP established in 2021^6–8^. TRIBE (Targets of RNA-binding proteins Identified By Editing) connects a RBP with the catalytic domain of adenosine deaminase ADAR (ADARcd) into a fusion protein^6^. When the RBP in the fusion protein binds to target RNAs, ADARcd edits adjacent adenosines (A) into inosines (I), which are read as guanosines (G) in high-throughput sequencing, thereby identifying the target RNAs of the RBP. In its improved version HyperTRIBE, ADARcd possesses an amino acid substitution so HyperTRIBE works with higher detection sensitivity and less bias in editing sites, that is, it requires lower sequencing depth and determines targets with lower false-negative rates compared with TRIBE^13–15^. TRIBE has been employed to flies^6,13,15–17^, mammals^13,18^, malaria parasite^19^ and planta^20^. RNA tagging^8^ is fused a RBP with the poly(U) polymerase (PUP-2)^21^ originated from *C. elegans* so it can add poly(U) tails to 3 ‘ ends of target RNAs and identify U-tailed mRNAs by paired-end sequencing in yeasts. As poly(U) polymerase modifies the 3’ ends of mRNAs, RNA tagging is supposed to work better for RBPs bound on 3’UTRs since the regions are close to 3’ ends. STAMP^7^ is designed to fuse the RNA cytosine deaminase APOBEC1^22^ from *Rattus norvegicus* with a RBP and to determine cytosine to uridine transitions in target mRNAs by sequencing in mammals. The recent research has indicated that STAMP works well in human cells but it is hardly applied in *Drosophila* cells^23^. Likewise, it is uncertain that RNA modification enzymes have activity in untested species and therefore utilization of these methods in other species needs further verification and optimization.

Yeast is remarkable eukaryotic model organisms for basic studies and also indispensable to bioengineering^24^. So far, approximately 120 yeast proteins are found to be associated with mRNAs^25^ and many of them are conserved in higher organisms. The typical RBP KHD1 is a *S. cerevisiae* homolog of mammalian heterogeneous nuclear ribonucleoprotein K (hnRNP K)^26^, and includes three conserved K homology (KH) domains. Under suitable environments, yeast cells undergo asexual reproduction mainly by budding. Genome screening has showed that KHD1 interacts with about 50% of bud localization mRNAs^27,28^, most of which encode membrane or secretory proteins^29^. In addition, KHD1 has been reported to have distinct effects on different target genes. For example, KHD1 inhibits translation of FLO11 mRNA^30^ while stabilizes MTL1 mRNA^28,31^. During asymmetric cell division^32–34^, KHD1 can participate in the translational inhibition of ASH1 mRNA and the accumulation of ASH1 protein in growing bud, thereby inhibiting mating type conversion in daughter cells^35^.

Another multi-functional protein BFR1 is involved in the uneven distribution of mRNA during yeast budding and responsible for target mRNA localization in bud tip^36^. It is localized to endoplasmic reticulum^25,37^ and is a component of the poly-ribosomal mRNA-binding protein complex^38^. BFR1, which was also found in the late P-body, is very important for the delayed entry of specific mRNA into the P-body, such as VNX1 and TDP1 mRNA^39^. It has been proposed that BFR1 protein may mediate the transformation of target mRNA from translation to degradation in cells under stress conditions such as starvation, resulted in the accumulation of BFR1 in P-body^39^.

Because the rigid cell wall leads to less efficiency of UV cross-linking for CLIP in organisms like plants, fungi, and bacteria, TRIBE could be a good choice in the cells with cell wall. In addition, multiple available tools for RBP target examination in one species will be conducive to identification of confident targets as well as to studying several cooperative RBPs in a large complex simultaneously. Thus, we adapted HyperTRIBE in fungus *S. cerevisiae* for the first time and investigated the target transcripts of RNA-binding proteins KHD1 and BFR1. Our results showed that HyperTRIBE performs well with low background, good sensitivity and reproducibility. Also, combining with differentially expressed genes (DEGs) after depletion of KHD1 or BFR1, we identified the direct targets of these RBPs and made implications for their functions. Our work established a simple and useful method in yeast and will be beneficial for the field.

## Results

### The KHD1-Hyper TRIBE successfully identified KHD1 targets

KHD1 protein possesses three KH domains, which are one of the well-known classical RNA-binding motifs (Table 1). KHD1 was reported to inhibit the translational initiation of hundreds of mRNAs in the process of mRNA transfer to specific cell loci, indicating that it has hundreds of potential mRNA targets^28^. To identify the targets of KHD1 in yeast, KHD1 was fused with the hyper-active catalytic domain of human ADAR2 E488Q (hADAR2cd) (Fig. S1A) because hADAR2cd had been used in mammalian HyperTRIBE^13^ and confirmed having activity in yeast cells^40^. The protein expression of hADAR2cd (Hyper only) and KHD1-hADAR2cd (KHD1 Hyper) in *S. cerevisiae* were validated by western blotting (Fig. S1B). Very few editing events were detected in the negative control Hyper-only, while 770 editing sites and 492 target transcripts were detected in KHD1-HyperTRIBE, indicating that HyperTRIBE works well in yeast cells (Fig. 1A).

**Table 1.**
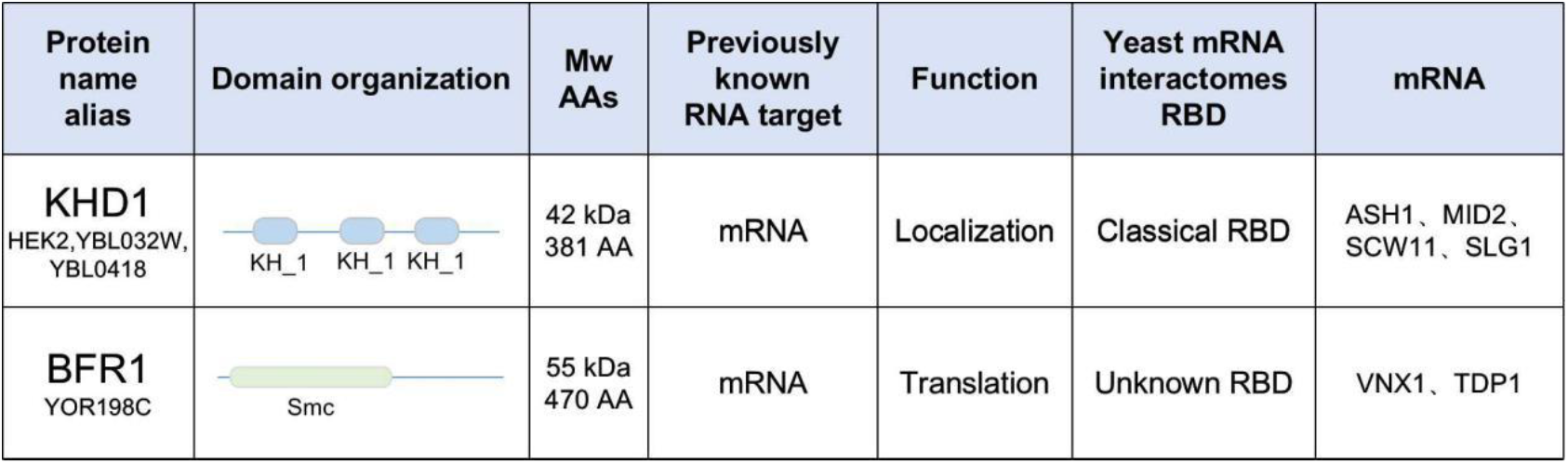
Information about yeast RNA-binding protein KHD1 and BFR1. KHD1 protein has three KH RNA-binding motifs, and has been confirmed to bind ASH1 mRNA and be necessary for the effective localization of target mRNAs^33^. BFR1 lacks typical RNA binding domains and participates in the protein secretion pathway^25,37^. It is located in the endoplasmic reticulum (ER) under normal conditions and in P body after stress^39^.

**Figure 1.**
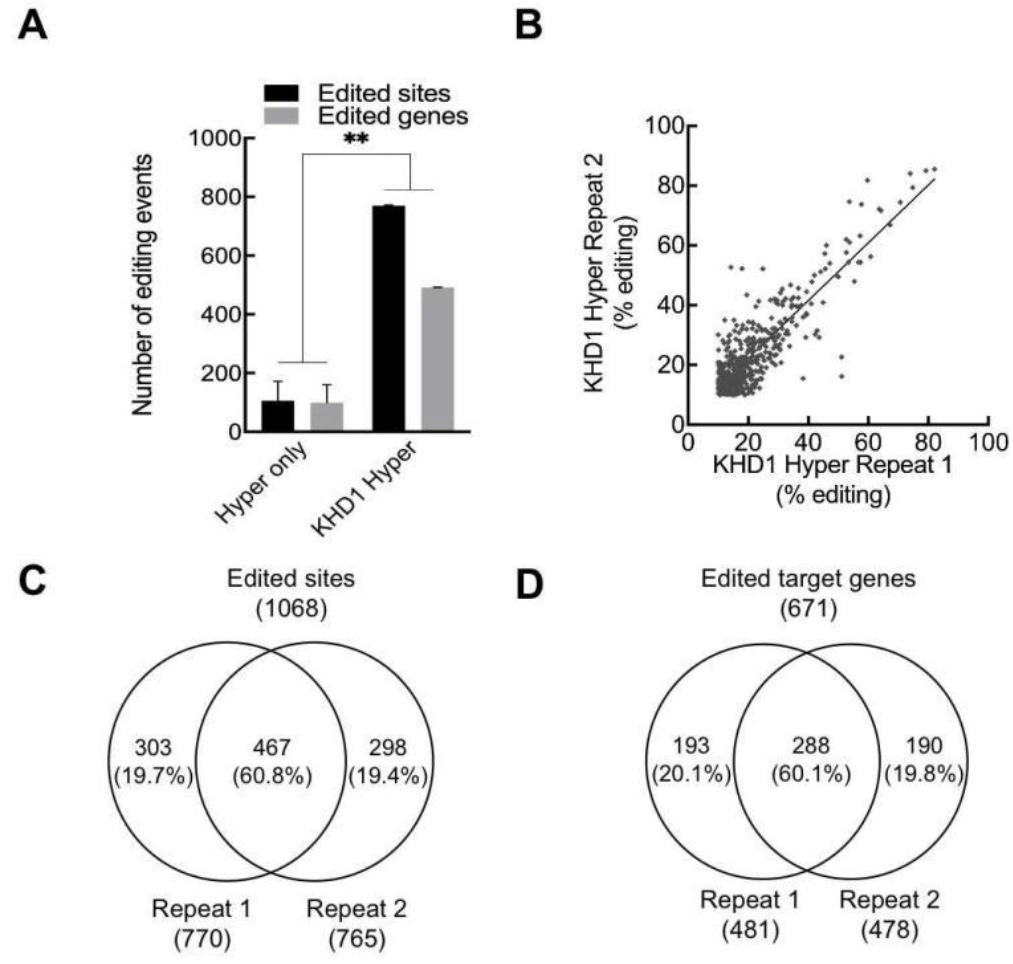
KHD1-HyperTRIBE reproducibly identifies target RNAs of KHD1 in yeast. **(A)** The editing sites and editing genes detected in KHD1-Hyper**TRIBE** are significantly higher than those detected in Hyper only. KHD1-HyperTRIBE detected **~**770 editing sites and **~**500 target genes (Editing≥10%, read≥20). *N* = 2, +SEM; \*\**P* < 0.01, paired one-tailed Student’s t test. **(B)** The editing degree of detected sites in two biological replicates is similar (R^2^=0.74). **(C, D)** About 60% of editing sites and editing genes are reproducibly detected in two replicates as showed in Venn diagram.

The editing sites were found in protein-coding regions and 3’ UTRs but not in 5’ UTRs (Fig. S2A), and these editing events in KHD1-HyperTRIBE were restricted to A-to-G nucleotide conversion, no other types of nucleotide conversion were significantly determined (Fig. S4). This observation indicated that these editing events were resulted from hADAR2cd activity not from non-specific or other kinds of enzyme activities. The editing events in KHD1-HyperTRIBE were highly reproducible both in frequencies and positions (Fig. 1B & 1C), revealing that the method specifically identifies target transcripts. With the threshold of read number 20 and editing percentage 10% (r20e10), 288 (60%) genes were marked in two biological replicates and defined as highly-confident targets (Fig. 1D). A total of 671 target genes was detected at least once in two experiments, taking up 11% of *S. cerevisiae* genes (Fig. 1D). When the threshold was set as read number 10 and editing percentage 10% (r10e10), a total of 835 targets was identified (Fig. 2E).

**Figure 2.**
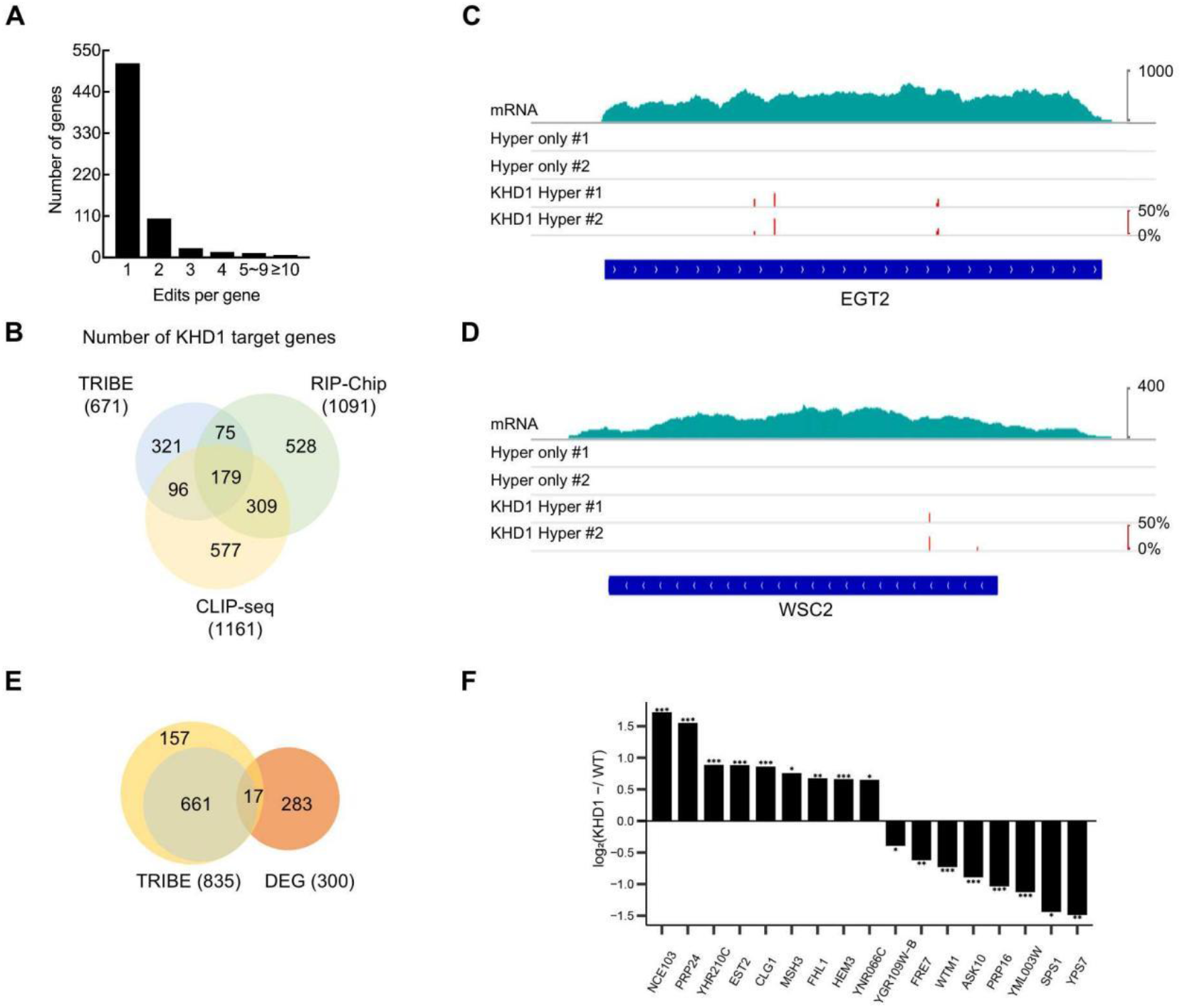
KHD1-HyperTRIBE targets overlap well with the KHD1 targets detected by other methods. **(A)** The frequency histogram of editing sites per gene shows that most target genes of KHD1 are edited once in KHD1-HyperTRIBE. **(B)** The comparison of targets detected in KHD1-HyperTRIBE, RIP-Chip (SGD), and CLIP-seq (POSTAR3) indicates that 350 (52%) of HyperTRIBE target genes have been detected in the other two methods. **(C, D)** The IGV view of well-known KHD1 targets shows that KHD1-HyperTRIBE specifically edits target RNAs with high editing efficiency. The editing sites and editing percentages of EGT2 and WSC2 are shown as short red bars, and the heights represent the editing percentages of loci. **(E, F)** Seventeen genes are overlapped between KHD1-HyperTRIBE targets and differentially expressed genes (DEG) in *khd1*-depleted (*khd1*^-^) yeasts observed by Microarray analysis (SGD). The threshold of yellow circle is r10e10, and the threshold of green circle is r20e10. The cutoff of DEG is *P*<0.1 and Log (*khd1*^-^/WT) value of ±0.2 to ±2 (E). For the overlapped 17 genes, the mRNA level changes in *khd1*^-^ yeasts compared with wild type (WT) are shown. The bar chart shows log_2_ fold-change value of mRNA expression (*khd1*^-^/WT). *N* = 3, \**P*<0.1, \*\**P*<0.05, \*\*\**P*<0.01, paired one-tailed Student’s *t* test (F). SGD: Saccharomyces Genome Database, IGV: Integrative Genomics Viewer, DEG: Differentially Expressed Genes.

The frequency histogram of editing sites per KHD1 target gene showed that 76% of target genes possess a single editing site, that is 24% has more than one editing sites (Fig. 2A). When the KHD1-HyperTRIBE target genes were compared with the results of RNA immunoprecipitation-Chip (RIP-Chip)^28^ and CLIP-seq^30^, half the HyperTRIBE targets was detected at least once in the other methods and this overlapping ratio was similar to that of the other two methods (Fig. 2B). Interestingly, the well-known bud tip localization mRNAs like MTL1, WSC2, EGT2 and IST2^29^ were identified as KHD1 targets in KHD1-HyperTRIBE (Fig. 2C, 2D & Table 2). In addition, 17 of KHD1-HyperTRIBE target mRNAs had significantly different expression in *khd1*-depleted yeasts compared with wild type yeasts (Fig. 2E & 2F), while up-regulation and down-regulation took similar proportions in these 17 genes (Fig. 2F), suggesting KHD1 binding possibly made different impacts on target mRNA levels.

**Table 2.** KHD1-HyperTRIBE identified 7 bud tip localization mRNAs as KHD1 targets. † In addition to ASH1, the other six transcripts are bud tip localization mRNA edited by KHD1-HyperTRIBE. The table summarized the reported information about 7 bud tip localization mRNAs^27,29^. Yes, 90% bud localization; Partial, 50–60% localization; Weak, 15–30% localization; No, unlocalized (5% localization); ND, not determined; -, not assayed. * IRC8 was not enriched in RIP^28^ but enriched in another set of RIP (q<0.05)^42^.

| Gene | RNA enriched in bud tip | Protein enriched in bud tip | Protein levels in KHD1 OE cells | RIP-Chip with KDH1 antibody | The colocalization of mRNA with KHD1 protein | RNA pulldown |
| --- | --- | --- | --- | --- | --- | --- |
| ASH1† | Yes | Yes | Down | ✓ | ✓ | ✓ |
| EGT2 | Partial | Yes | - | ✓ | ✓ | - |
| IST2 | Yes | Yes | ND | ✓ | ✓ | - |
| WSC2 | Yes | Yes | ND | ✓ | ✓ | - |
| TAM41 | Yes | No | - | X | X | - |
| IRC8 | Yes | Yes | - | X* | - | - |
| MTL1 | Weak | ND | UP | ✓ | ✓ | ✓ |

To find out the consensus sequence in the target mRNAs, we carried out MEME motif analysis using the regions including 100 bp upstream and 100 bp downstream from the TRIBE editing sites. The analysis showed that YCAACAA and YCAUCAU (Y represents a pyrimidine, U or C) motifs were enriched in KHD1 binding regions (Fig. 3A), consistent with structural studies of Nova third KH domain which specifically interacts with the internal CA in a YCAY (Y represents a pyrimidine, U or C) motif^41^ and similar to previously published CLIP-seq data^30^. To explore whether the binding of KHD1 protein on its target transcripts affects target mRNA levels, the steady state mRNA abundance changes after knocking out *khd1* were compared among three different groups of genes, KHD1-TRIBE targets, KHD1-TRIBE targets with above motif (YCAACAA), and non-targets (Fig. S2C). Cumulative frequency distributions of the mRNA level changes showed that KHD1-TRIBE targets with specific motif had slightly decreased distributions compared with the other two groups, indicating KHD1 stabilized its target mRNAs with this motif (Fig. S2C).

**Figure 3.**
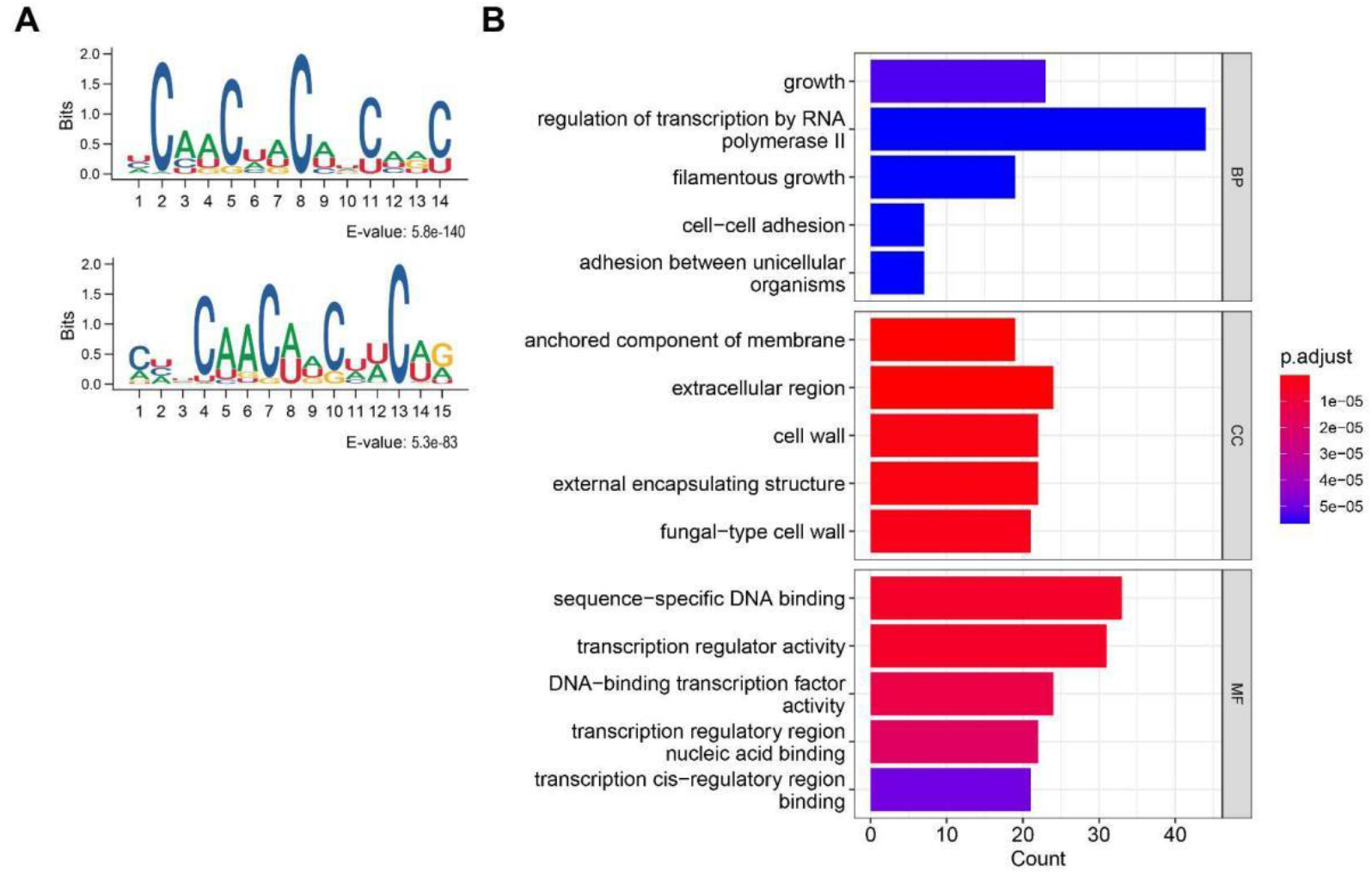
The targets of KHD1-HyperTRIBE have specific motifs and their functions are enriched in filamentous growth and cell adhesion. **(A)** The motifs identified by KHD1-HyperTRIBE include CAACAA and CAUCAU, which are consistent with the results of KH domain structure and KHD1 CLIP-seq analysis published. **(B)** GO term analysis indicates that the roles of KHD1 target genes are enriched in transcriptional regulation, cell growth, filamentous growth, and cell adhesion, which correlate well with reported *khd1*-knockout phenotypes. BP: Biological Process, CC: Cell Composition, MF: Molecular Function.

Gene ontology (GO) term analysis showed that target genes of KHD1 are related to filamentous growth, cell growth and adhesion (Fig. 3B). Interestingly, it was reported that the *khd1*-knockout yeasts have increased filamentous growth, ability of adhere together and to agar plate^30^, indicating that KHD1-HyperTRIBE correctly captured KHD1 targets. It also suggested that the binding of KHD1 to its target transcripts controls the target expression in certain ways thereby inhibiting filamentous growth and cell adhesion. TRIBE results provide a basis for further functional and mechanism study of KHD1.

### The BFR1-HyperTRIBE exhibits BFR1-determined editing activity

To establish hyperTRIBE in yeast, we further tested the method by using another multi-functional RBP BFR1. Even though BFR1 includes only a Structural Maintenance of Chromosomes (Smc) domain and lacks any classical RNA-binding domains (Table 1), it was well known to be widely involved in mRNA metabolism. The results showed that in *Saccharomyces cerevisiae* cells, the proteins of hADAR2cd (Hyper-only) and BFR1-hADAR2cd (BFR1-Hyper) were expressed well after induction. Then, the numbers of edited sites and edited genes were counted in Hyper-only and BFR1-Hyper RNA-sequencing results. The 106 edited sites and 100 edited genes were detected in the negative control Hyper-only group, while 847 edited sites and 700 edited genes were observed in BFR1-Hyper group, the ratio of signal to noise is more than seven times (Fig. 4A). The high level of A-to-G editing but no other types of editing was observed in BFR1-Hyper samples (Fig. S4). The editing sites were observed in protein-coding region and 3’ UTR but not in 5’ UTR (Fig. S2B). In addition, the editing events were repeatable in their editing percentages, positions and target genes (Fig. 4B-4D), and ~61% of target genes were reproducibly detected (Fig. 4D), indicating that the target identification in BFR1-HyperTRIBE was highly specific.

**Figure 4.**
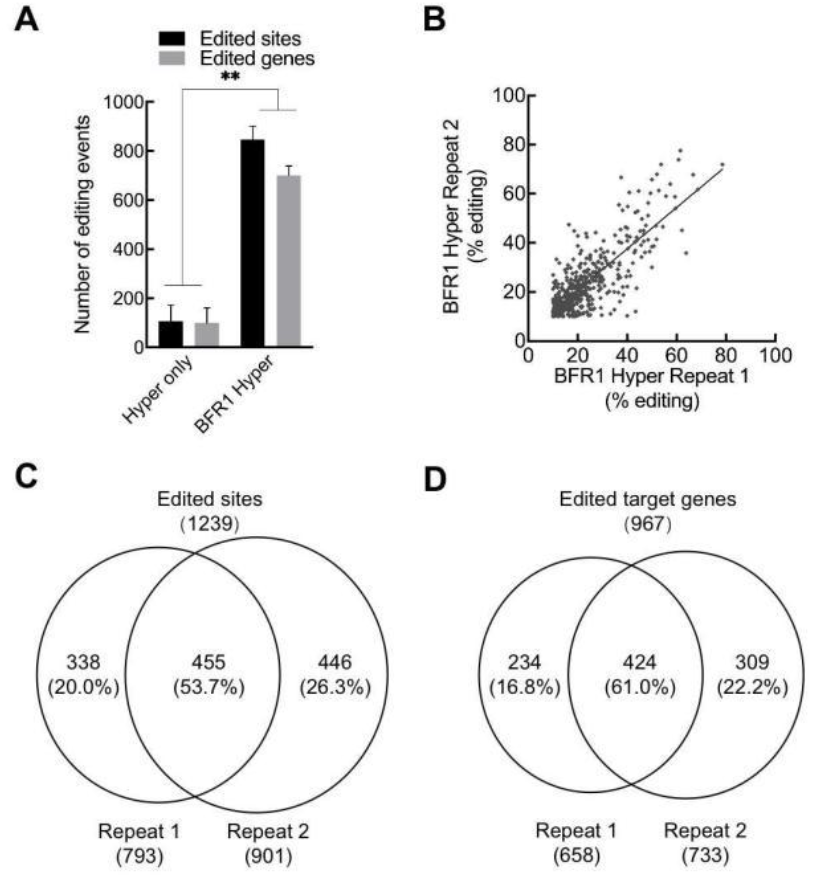
BFR1-HyperTRIBE successfully edits the target RNAs of BFR1p with high reproducibility. **(A)** BFR1-HyperTRIBE generates much more edited sites and edited genes than Hyper-only, indicating BFR1-HyperTRIBE efficiently identifies BFR1 targets (Read≥20, editing≥10%). *N* = 2, +SEM; \*\**P* < 0.01, paired one-tailed Student’s t test. **(B)** The editing percentages of detected sites in two biological repeats are similar (R^2^=0.63). **(C, D)** Venn diagram shows that ~60% of editing sites and editing genes are reproducibly detected in two replicates.

The frequency histogram of editing sites per BFR1 target gene showed that most genes have only one editing site (Fig. 5A). When a total of 967 target genes (r20e10), which was detected at least once in two BFR1-HyperTRIBE experiments, was compared with the published BFR1-RNA Tagging data^8^ and RIP-Chip data^42^, about half of BFR1-HyperTRIBE targets were detected at least once in the other two methods (Fig. 5B). The representative BFR1-HyperTRIBE target genes like SCEC27 and OLA1 were identified in two independent experiments with high editing percentages (Fig. 5C & 5D), and SCEC27 was known to be involved in the transport between endoplasmic reticulum and Golgi apparatus^43^ while OLA1 might regulate mRNA translation^44^.

**Figure 5.**
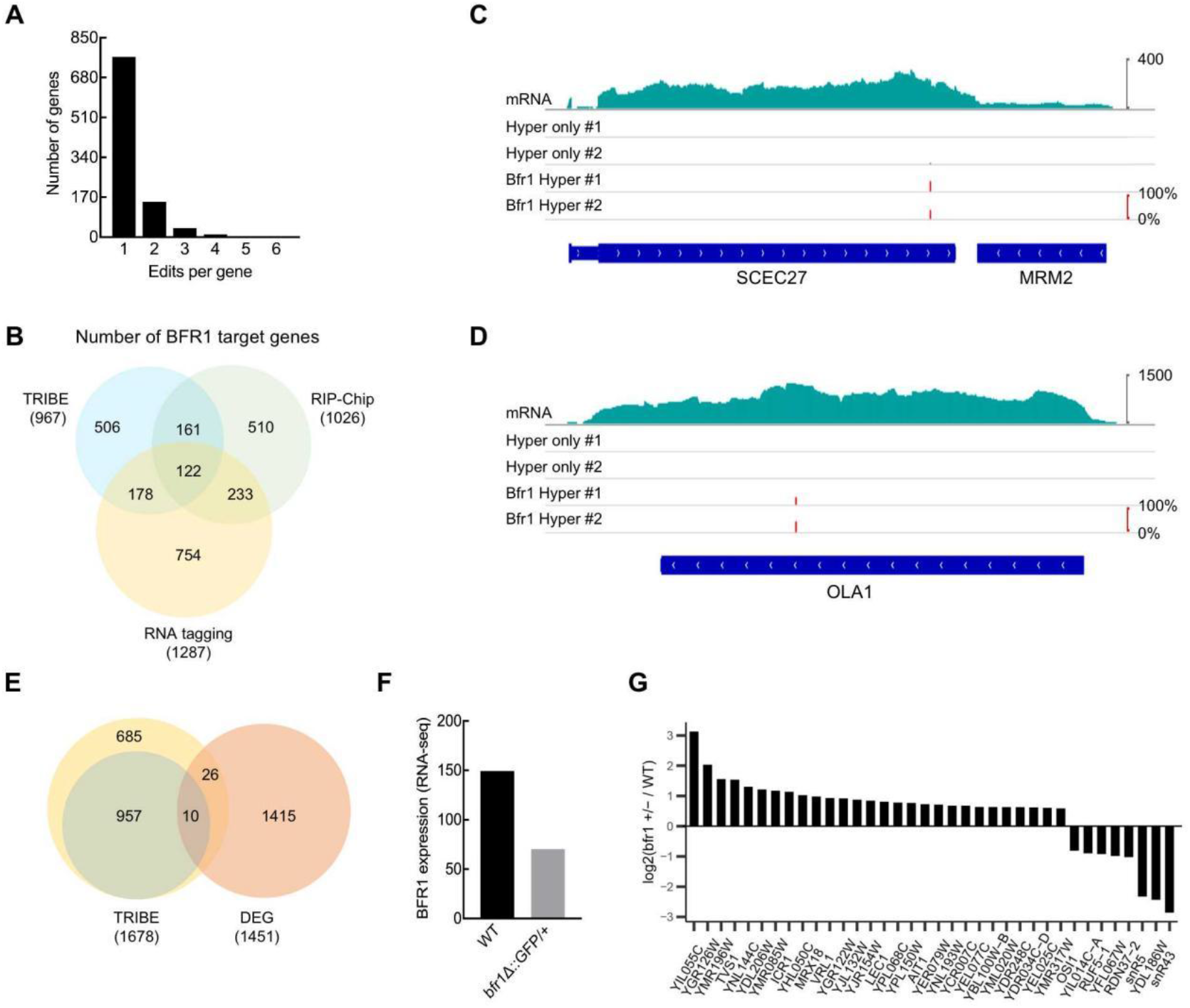
BFR1-HyperTRIBE targets overlap well with the BFR1 targets detected by other methods. **(A)** The frequency histogram of editing sites per gene shows that most target genes of BFR1 are edited once in BFR1-HyperTRIBE. **(B)** The comparison of targets detected in BFR1-HyperTRIBE, RIP-Chip^42^, and RNA tagging^8^ indicates that half of HyperTRIBE target genes have been detected in the other two methods. **(C, D)** The IGV view of two representative BFR1 targets detected in BFR1-HyperTRIBE. The editing sites and editing percentages of SCEC27 and OLA1 are shown as short red bars, and the heights represent the editing percentages of loci. **(E-G)** Thirty six genes are overlapped between BFR1-HyperTRIBE and DEG in *bfr1*^+/-^ yeasts observed by RNA-seq (SGD). The cutoff of DEG is FPKM≥2 and FPKM ratio multiples are greater than 1.5. The TRIBE threshold is r10e10 for yellow circle and r20e10 for green circle (E). The BFR1 mRNA expression in the *bfr1^+/-^* yeasts is about half of WT (F). The bar chart shows log_2_ mRNA-level-fold-change-value of overlapped 36 genes (*bfr1*^+/-^/WT) (G). SGD: Saccharomyces Genome Database, DEG: Differentially Expressed Genes.

Furthermore, 36 overlapping genes discovered in 1678 BFR1-HyperTRIBE target genes (r10e10) and differentially expressed genes from RNA-Seq data between *bfr1^+/-^* and WT^45^ (Fig. 5E). The *bfr1^+/-^* strain had lower BFR1 mRNA level compared with wild type strain (Fig. 5F). Thus, the partial depletion of BFR1 resulted in transcriptome change of above 36 overlapped genes, including 28 increased and 8 decreased genes (Fig. 5G). Cumulative frequency distributions of the mRNA abundance changes after partial depletion of BFR1 expression were compared between two groups, BFR1-TRIBE target group and non-target group (Fig. S2D). The result showed a slight but significant difference between two groups, suggesting that BFR1 mainly destabilizes the target mRNAs (Fig. S2D). These observations were consistent with the reported research that BFR1 could trigger mRNA degradation under stress conditions like starvation^39^.

In addition, BFR1 was known to be localized to endoplasmic reticulum (ER)^25,37^ so we examined whether BFR1 targets were enriched for ER-translated mRNAs. The cumulative frequency distributions of mRNA enrichment on ER showed that BFR1-TRIBE targets were significantly enriched for abundant, ER-translated mRNAs in comparison to all mRNAs or non-targets (Fig. S3A). These targets were similarly enriched for both SEC complex-dependent and SEC complex-independent translocation events (Fig. S3B, S3C). GO term analysis of BFR1 target genes showed that most of its target genes were involved in metabolic processes and the membranous organelle-related pathways typically ER membrane (Fig. 6), which is in agreement with the previous report^38^.

**Figure 6.**
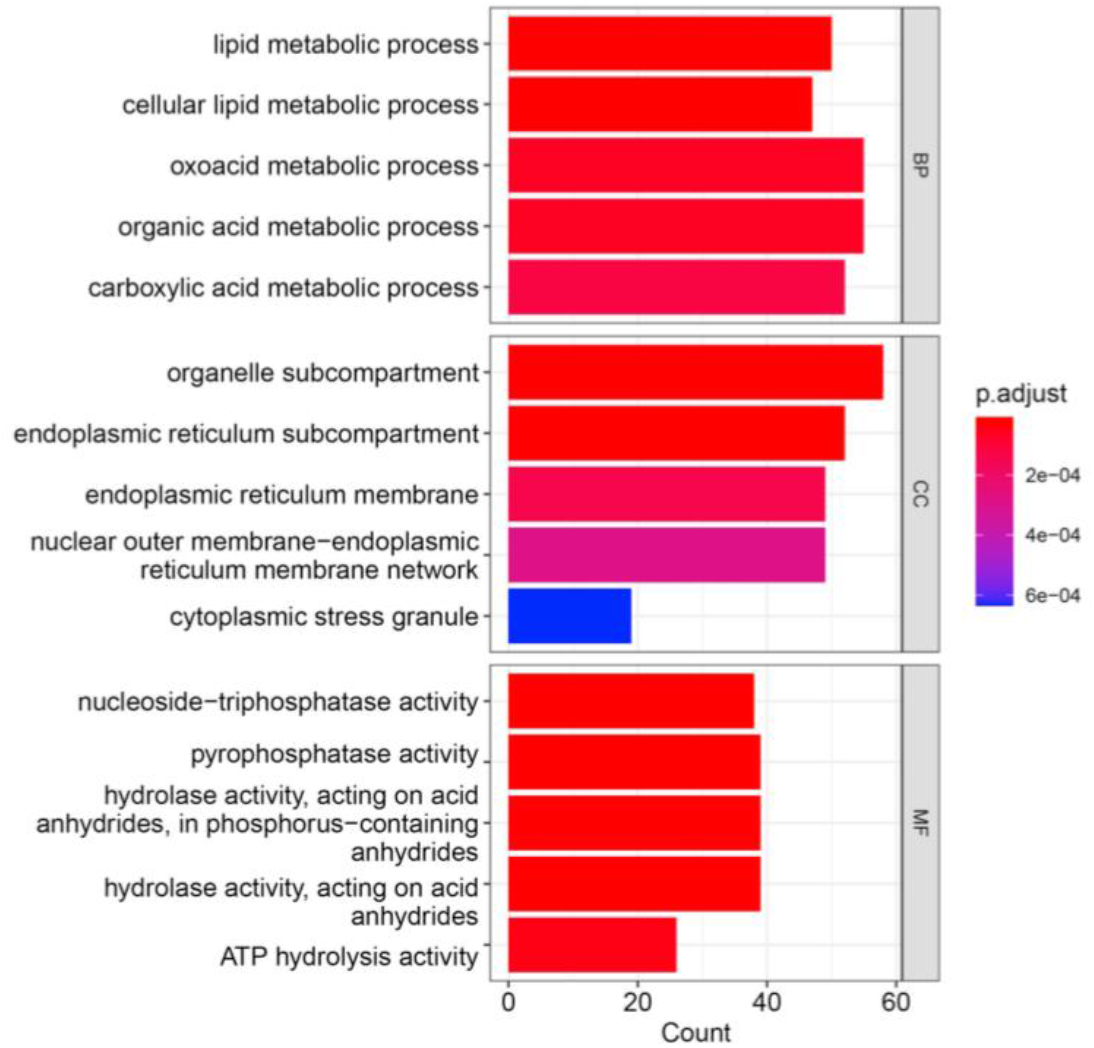
BFR1-TRIBE targets are highly enriched in metabolic processes and membranous organelle-related pathways. The GO term analysis results of biological process (BP), cell composition (CC), and molecular function (MF) are shown.

## Discussion

Our HyperTRIBE data of KHD1 and BFR1 are well correlated with their published functions. The roles of KHD1 targets are enriched in the processes of filamentous growth, cell growth and adhesion (Fig. 3B), thus, this part of targets should be responsible for the phenotype of *khd1*-knockout yeast like increased filamentous growth, ability of adhere together and to agar plate^30^. Our data also suggest that KHD1 stabilizes its targets with specific motif (Fig. S2C). BFR1 HyperTRIBE targets are enriched with mRNAs translated on ER (Fig. 6 & Fig. S3) and BFR1 more likely destabilizes its targets (Fig. S2D).

As master regulators in cells, RBPs take part in gene regulation at multiple levels by specifically recognizing RNA. To reveal the underlying mechanisms, the methods profiling RBP-RNA interaction network have been rapidly developed. These methods have their own advantages. HITS-CLIP and variants can detect the RNA targets from endogenous RBPs with relatively accurate binding site information. Instead, TRIBE, RNA tagging and STAMP are antibody-free methods so can avoid the problems arising from the limited specificity of antibody. Also, they can be the alternative choices for discovering RBP targets in cells with cell wall structures such as plants, fungi and bacteria, where the low UV-crosslinking efficiency might be a problem. In particular, TRIBE and STAMP need a small number of starting cells and undergo a regular RNA-seq library preparation procedure so they are easier to operate and can be applied to the studies at a single-cell level. In addition, the research on a complex comprising multiple RBPs will benefit from combining different techniques in the same organism, for example, different RBPs, fused with different RNA modification enzymes, can be expressed into the same cells to mark the common and unique targets of the RBPs. Thus, the development of varying strategies profiling the RBP-RNA interactions will greatly improve the research on RNA biology.

The fusion protein-based system has comprised an important part in RBP target identification methodology. The key point for establishment of this kind of method in an untested species is to seek out a RNA modification enzyme with high activity in the species. The development of TRIBE in *Drosophila* used the fusion protein of *Drosophila* ADARcd (dADARcd) with a RBP^6^ and improved version HyperTRIBE employed hyper-active dADARcd (E488Q)^13–15^. In mammalian HyperTRIBE, hADAR2cd (E488Q) was used to construct fusion protein due to higher editing efficiency of hADAR2cd (E488Q) than dADARcd (E488Q) there^13^. To adapt HyperTRIBE in yeasts, we tried the hyper-active catalytic domain from dADAR (E488Q), human ADAR2 (E488Q), and human ADAR2 (E488Q, V493T, N597K)^40^ in constructing TRIBE fusion proteins. The HyperTRIBE with dADAR (E488Q) or hADAR2 (E488Q, V493T, N597K) failed to produce results with consistently high signal to noise ratio (Fig S5 and data not shown). Instead, the HyperTRIBE with hADAR2 (E488Q) was working well with a high signal to noise ratio, great reproducibility and remarkable consistency with published data of tested RBPs. Our work established a simple and useful HyperTRIBE method in yeast for the first time and will be beneficial for the field.

## Materials and Methods

### Plasmid construction

RBPs of interest were cloned into pcDNA3-3HA-hADAR2cd-E488Q^13^ to build pcDNA3-3HA-RBP-hADAR2cd-E488Q using Gibson assembly method^46^ (EasyGeno Single Assembly Cloning kit, Tiangen). PGK1 and CYC1 were acquired from yeast gDNA. PGK1, 3HA-RBP-hADAR2cd-E488Q and CYC1 were cloned into pRS313 to get pRS313-PGK1-3HA-RBP-hADAR2cd-E488Q-CYC1. The V493T/N597K mutations were introduced into hADAR2cd-E488Q by using Fast Site-Directed Mutagenesis Kit (Tiangen). Myc tag substituted Khd1 to produce pRS313-PGK1-hADAR2cd-E488Q-CYC1 plasmid. PGK1 was replaced by the PCR product of GAL1 to make inducible plasmids. All cloned sequences were verified by Sanger sequencing.

### Yeast transformation

All *Saccharomyces cerevisiae* strains were generated from BY4742 (*MATα; his3Δ1; leu2Δ0; lys2Δ0; ura3Δ0*). The transformation method was adapted from prior protocol^47^. Centrifugation was completed for 5 min at 3000 rcf at room temperature (RT) unless otherwise specified. PEG 3350 and LiAc were autoclaved prior to use. A fresh streaked colony of BY4742 was inoculated in a 2-5 mL YPD medium (yeast extract, peptone, 2% autoclaved and isolated dextrose) at 30°C and 240 rpm for 12-16 h. Then, cultured yeasts were added into a fresh YPD medium at OD_600_ ~0.2, and the mixture was shaked until OD_600_ reached 0.6-0.7. The Pellet from 1.25 mL cultivation was washed once with 1 mL sterile water then immersed in 100 μL 1 M LiAc for 5 min at RT. The supernatant was removed by centrifuging for 3 min at 3000 rcf at RT. 33% PEG 3350 (w/v, mixed by pipetting immediately after added), 100 mM LiAc, 0.28 mg/mL salmon sperm DNA (pretreatment with 95°C heated for 5 min and ice-cold instantly), and 0.5-2 μg plasmid were added to remains in order, pipetting sterile water up to total volume of 360 μL, blending gently for 1 min. The compound was incubated at 30°C for 30 min and heat shocked at 42°C (for BY4742) for 20-30 min. The pellet was collected after centrifuging for 3 min at RT. 1 mL YPD was added for growing another 2 h at 30°C and 150 rpm. Cells were cleaned with sterilized water twice. The resuspended cells were plated on SD/-His (2% dextrose) agar and cultured for 2-4 days.

### Yeast cell cultures for TRIBE experiments

For the plasmids with PGK1 promoter, the transfected cells were grown in SD/-His media for 12~16 h. For the plasmids with GAL1 promoter, selected cells were maintained in SD/-His first, then saturated cultures were seeded into SD/-His media at OD_600_<0.2. When the OD_600_ was close to 1, uninduced control cells were collected for Western Blot (WB). Glucose was eliminated by resuspending yeasts three times in SC/-His media (without dextrose) as it could inhibit GAL1-initiated transcription^48^. Incubation was continued in SG/-His (with 2% galactose) media at OD_600_ = 1 for several hours. The expression of TRIBE constructs was examined by WB and the verified samples were used for RNA preparation.

### Purification of gDNA and RNA

Yeast cells were centrifugated at 6,000 rcf at 4°C. Cordless motor and pellet pestles (Fisher Scientific™) were utilized for cell homogenization in 100-200 uL of TRIzol^™^ reagent (Thermo Fisher Scientific). Grinding for 30 s and chilling on ice for 30 s were repeated five times for cell homogenization. Then, TRIzol^™^ reagent was further added to final volume of 1 mL per 50-70 μL of yeast pellet. Total RNA purification was followed the manufacturer’s instructions.

Extraction of gDNA was carried out in agreement with *Molecular Cloning: A Laboratory Manual^49^*. Acid-washed beads were prepared by steeping overnight in acid solution (6M HCl:H2O=1:50). After washing with ultrapure water ten times, beads were autoclaved and dried.

### TRIBE RNA-seq library preparation and analysis

Preparation of TRIBE RNA-sequencing library was followed the manufacturer’s instructions of Illumina Stranded mRNA Prep in which dUTP was included during second strand cDNA synthesis. UDI (IDT for Illumina UD Indexes) was functioned in sample pooling. The quality of NGS library was monitored by Qubit, dsDNA 915 kit for Fragment Analyzer^™^ (Agilent 5400)^50^, and qPCR (quantification of adaptor-added fragment). The qualified samples were sequenced by a paired-end–sequencing strategy (PE150) applying NovaSeqTM6000 v1.5 reagent kit on Illumina NovaSeq 6000 system. Two independent replicates of HyperTRIBE, KHD1-HyperTRIBE, and BFR1-HyperTRIBE experiments were carried out.

Raw data were processed as the published protocol^14^ with modification. Briefly, Cutadapt version 4.1 was used for adaptor dislodgment. Trimmomatic (0.38) was changed to paired-end sequencing model for inferior read filtering. Clean data were aligned to Saccharomyces cerevisiae genome (version R64-1-1) from SGD by STAR2.5.3a (--outFilterMismatchNoverLmax 0.10 --outFilterMatchNmin 16 --outFilterMultimapNmax 1), while alignment index was generated from Saccharomyces_cerevisiae.R64-1-1.dna.toplevel.fa and Saccharomyces_cerevisiae.R64-1-1.103.gtf. After removing PCR duplicates, editing events were identified with the indicated threshold of reads and editing percentages and single-nucleotide polymorphisms were avoided by wtRNA-RNA approach.

### Database

RNA IP-Chip data of KHD1 were from Saccharomyces Genome Database (https://www.yeastgenome.org/reference/S000127887). KHD1 targets detected in CLIP-seq were obtained from the database of POSTAR3 CLIPdb Module. Differentially expressed genes (DEG) of KHD1 were calculated from microarray data of wild type strain S288C and *khd1*-knockout strain *hek2Δ* from NCBI GEO Sample GSM843507 (https://www.ncbi.nlm.nih.gov/geo/query/acc.cgi?acc=GSM843507).

The RNA-tagging data of BFR1 were from SGD (https://www.yeastgenome.org/reference/S000182016), and the RIP-Chip data of BFR1 were from SGD (https://www.yeastgenome.org/reference/S000128183).

Differentially expressed genes (DEG) of BFR1 were calculated comparing RNA-Seq data of wild type strain BY4743 with BFR1 heterozygote mutant *BFR1*/*bfr1Δ::GFP* (*bfr1*+/-) obtained from NCBI GEO Sample GSM4240918 (https://www.ncbi.nlm.nih.gov/geo/query/acc.cgi?acc=GSM4240918).

### Motif analysis and GO term analysis

Online motif analysis was achieved by MEME version 5.4.1^51^. GO enrichment was operated in the R Programming Language. ID type of GENENAME was converted to ENTREZID with function bitr loaded with org.Sc.sgd.db (yeast genome ID) before proceeding to enrichGO^52^ function.

## Acknowledgments

We thank Yixin Huo for pRS313 plasmid, GAL1-containing plasmid and BY4742 yeast strain. The work was supported by the General Program of National Natural Science Foundation of China (31970622).

## Supplemental Materials

**Supplementary Fig 1.**
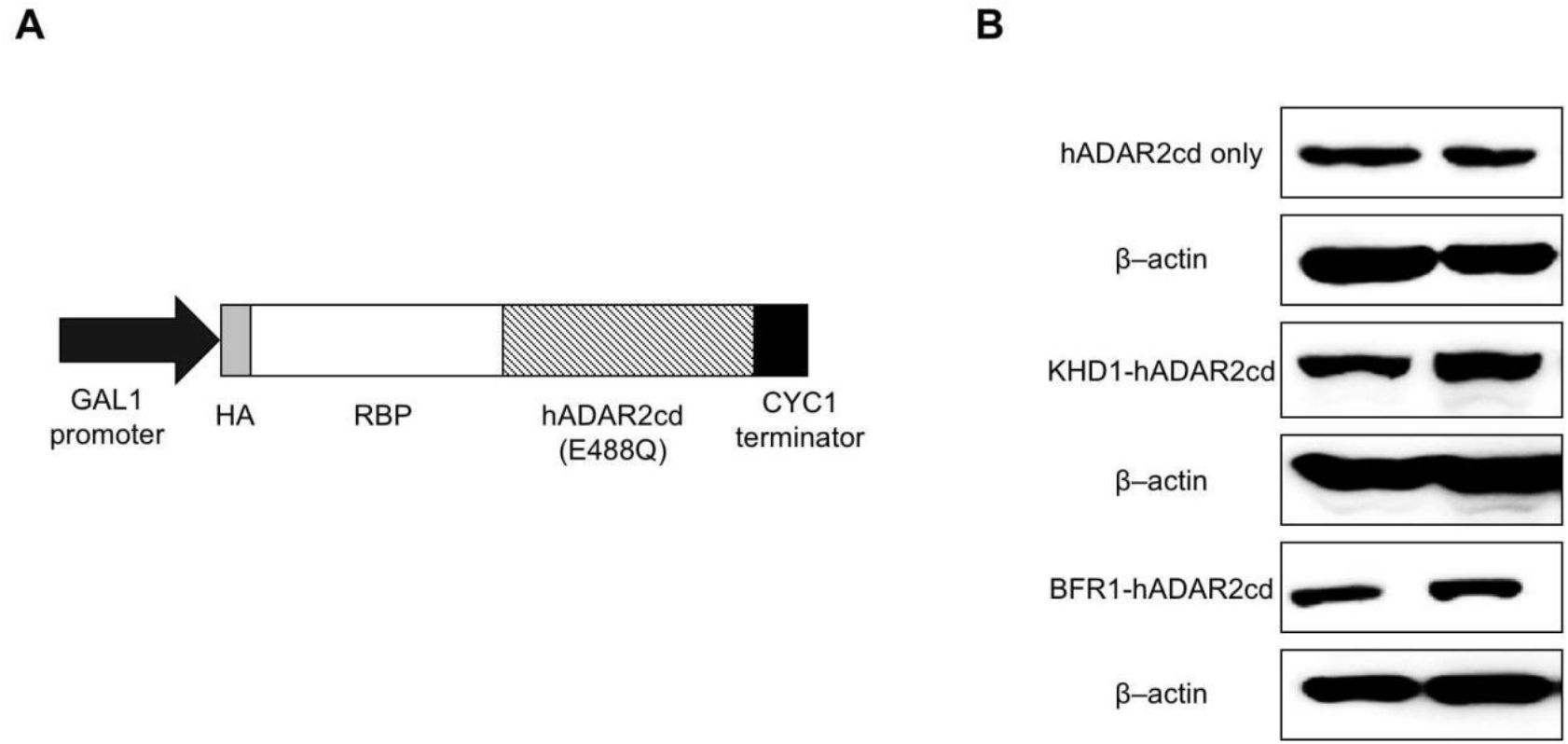
The TRIBE fusion protein expression is confirmed by Western blotting. **(A)** The map of DNA construct to express the HyperTRIBE fusion protein. GAL1 is a galactose inducible promoter, and hADAR2cd (E488Q) is the catalytic domain of human ADAR2 with a point mutation E488Q. **(B)** Western blot analysis shows the expression of HyperTRIBE fusion proteins in yeast. The hADAR2cd, KHD1-hADAR2cd, and BFR1-hADAR2cd were detected using the antibody against HA-tag. *β*-actin was used as a loading control.

**Supplementary Fig 2.**
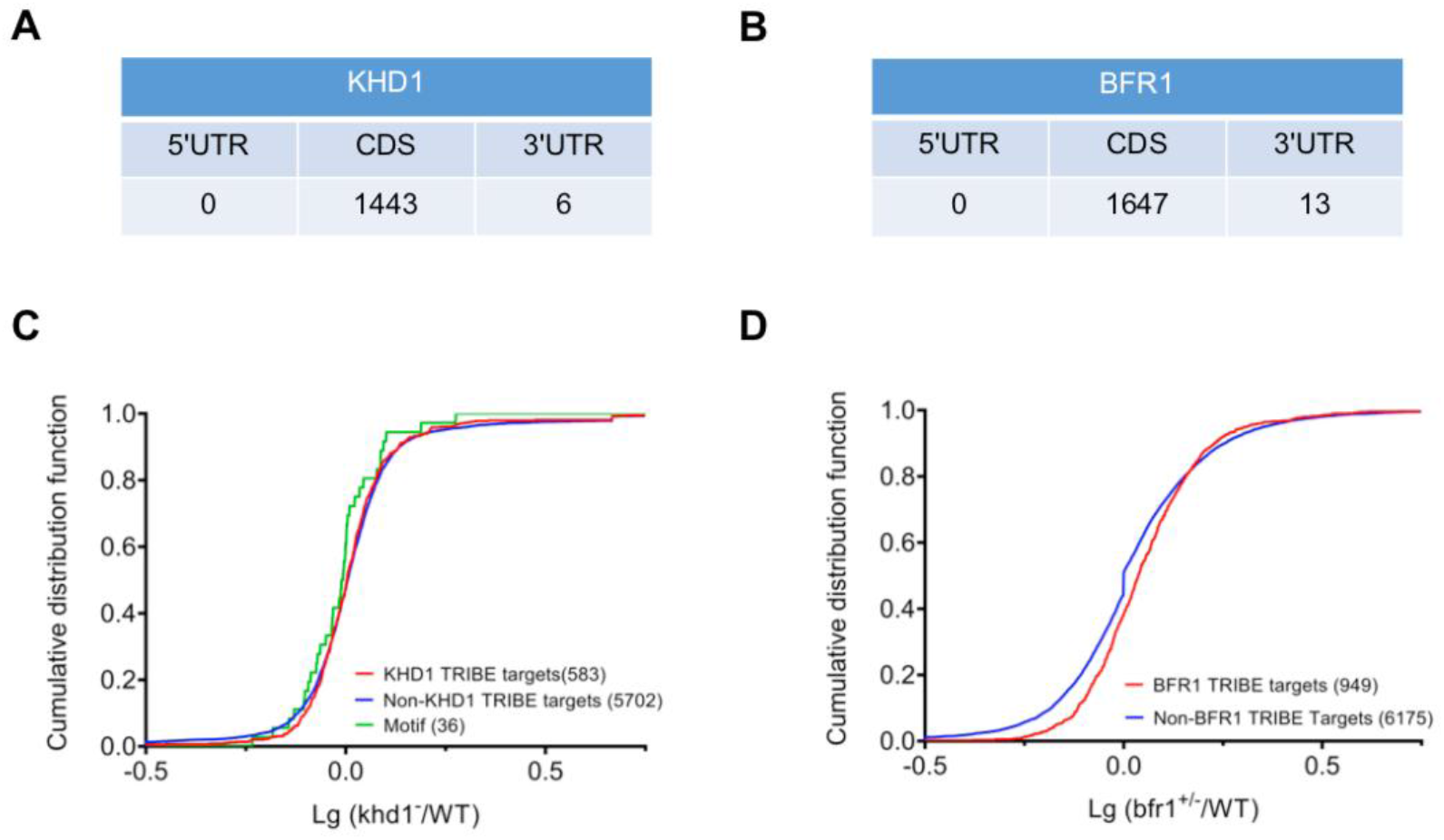
KHD1 stabilizes its targets with specific motif and BFR1 more likely destabilizes its targets. **(A, B)** The editing sites of KHD1-HyperTRIBE (A) and BFR1-HyperTRIBE (B) are mainly located in the protein coding region. **(C)** Cumulative distributions of log_10_ mRNA-abundance-ratio of *khd1* knockout yeast (*khd1^-^*) to its wild type counterpart (WT) from KHD1-TRIBE target group (red line, n=583), non-target group (blue line, n=5702) and motif (YCAACAA)-included target group (green line, n=36). **(D)** Cumulative distributions of log_10_ mRNA-abundance-ratio of *bfr1* heterozygote yeast (*bfr*^+/-^) to its wild type counterpart (WT) from BFR1-TRIBE target group (red line, n=949) and non-target group (blue line, n =6175).

**Supplementary Fig 3.**
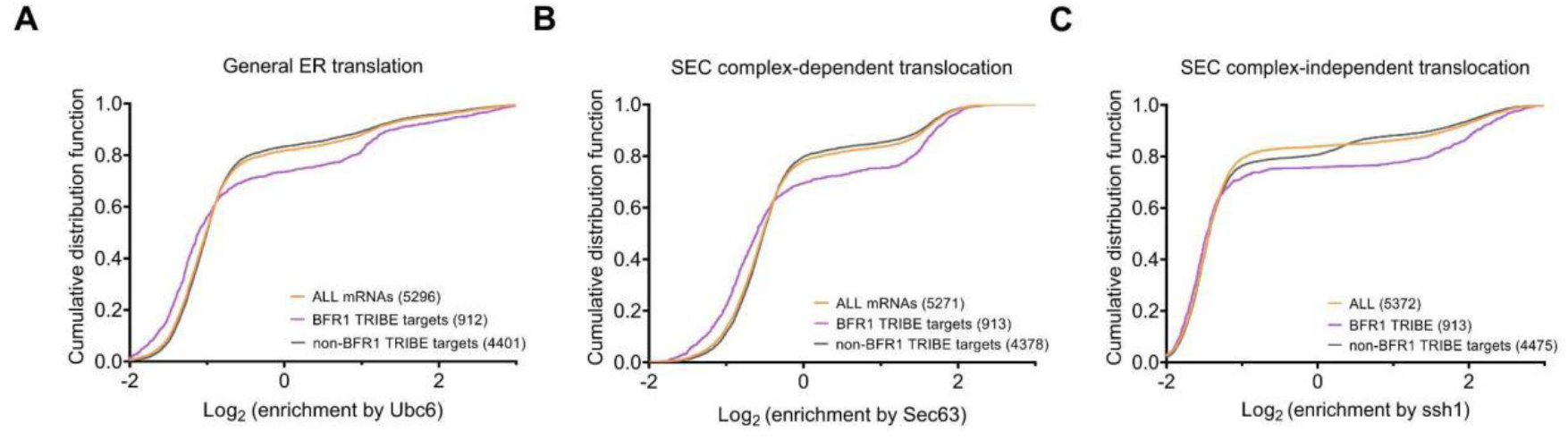
BFR1 targets are enriched with mRNAs translated on ER. Cumulative distributions are plotted for the groups of BFR1-HyperTRIBE targets, non-targets, and all expressed mRNAs for the following attributes: enrichment for mRNAs bound by ribosomes at the ER (log2(ubc6.7mchx enrichment)) (A), at the SEC complex (log2(sec63.7mchx enrichment)) (B), and at the SSH1 translocon complex (log2(ssh1.heh2.7mchx enrichment)) (C), obtained from the published ER-specific ribosome profiling experiments^53^.

**Supplementary Fig 4.**
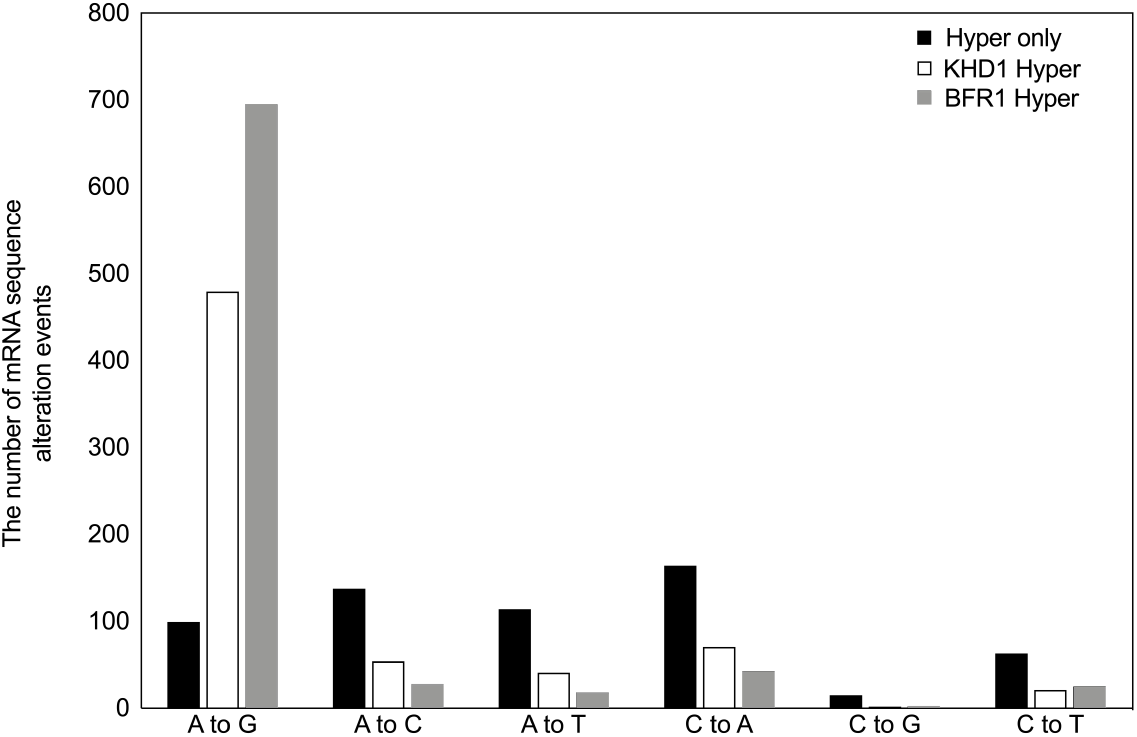
The mRNA sequence alterations detected in HyperTRIBE-expressed cells relative to wild type cells. Only A-to-G transitions are frequent in KHD1-HyperTRIBE and BFR1-HyperTRIBE samples but not in Hyper-only samples (N = 2, r20e10).

**Supplementary Fig 5.**
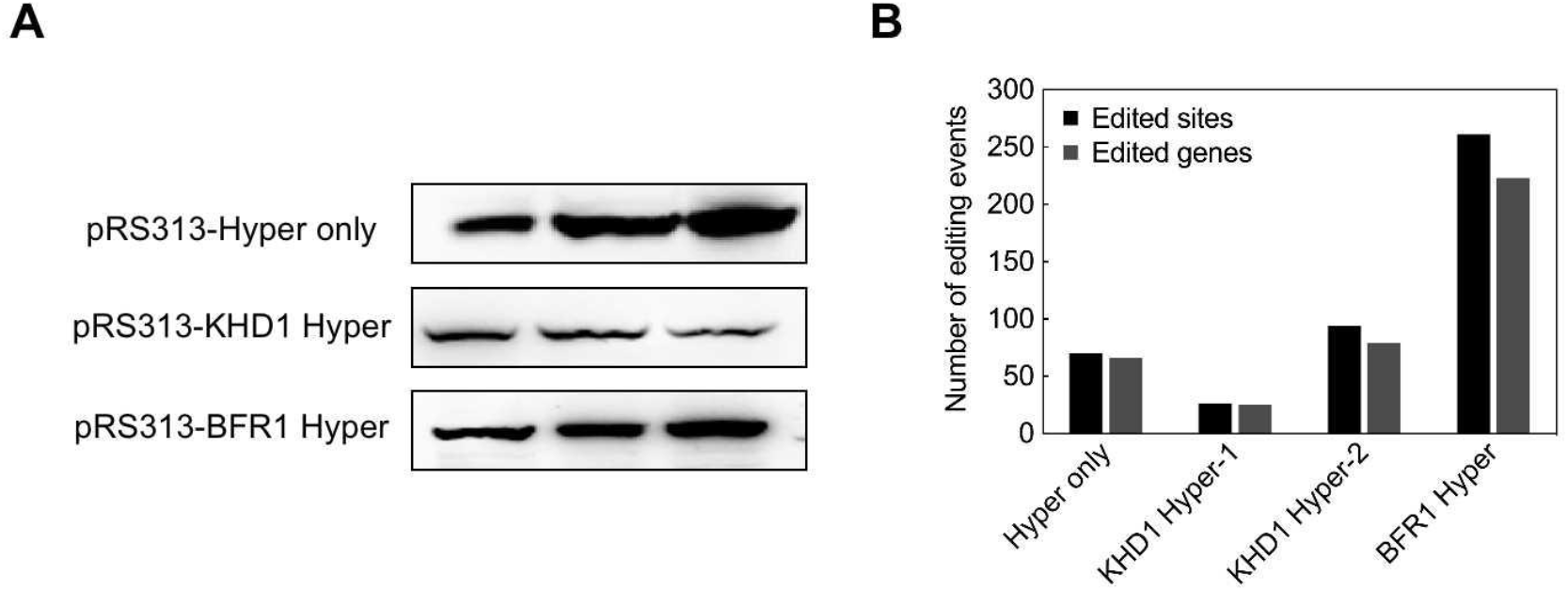
HyperTRIBE using hADAR2cd with three point mutations (E488Q, V493T, N597K) didn’t work consistently.

## References

1 Gerstberger, S., Hafner, M. & Tuschl, T. A census of human RNA-binding proteins. Nat Rev Genet 15, 829–845, doi: 10.1038/nrg3813 (2014).

2 Baltz, A. G. et al. The mRNA-bound proteome and its global occupancy profile on protein-coding transcripts. Mol Cell 46, 674–690, doi: 10.1016/j.molcel.2012.05.021 (2012).

3 Castello, A. et al. Insights into RNA biology from an atlas of mammalian mRNA-binding proteins. Cell 149, 1393–1406, doi: 10.1016/j.cell.2012.04.031 (2012).

4 Ramanathan, M., Porter, D. F. & Khavari, P. A. Methods to study RNA-protein interactions. Nat Methods 16, 225–234, doi: 10.1038/s41592-019-0330-1 (2019).

5 Van Nostrand, E. L. et al. A large-scale binding and functional map of human RNA-binding proteins. Nature 583, 711–719, doi: 10.1038/s41586-020-2077-3 (2020).

6 McMahon, A. C. et al. TRIBE: Hijacking an RNA-Editing Enzyme to Identify Cell-Specific Targets of RNA-Binding Proteins. Cell 165, 742–753, doi: 10.1016/j.cell.2016.03.007 (2016).

7 Brannan, K. W. et al. Robust single-cell discovery of RNA targets of RNA-binding proteins and ribosomes. Nat Methods 18, 507–519, doi: 10.1038/s41592-021-01128-0 (2021).

8 Lapointe, C. P., Wilinski, D., Saunders, H. A. & Wickens, M. Protein-RNA networks revealed through covalent RNA marks. Nat Methods 12, 1163–1170, doi: 10.1038/nmeth.3651 (2015).

9 Licatalosi, D. D. et al. HITS-CLIP yields genome-wide insights into brain alternative RNA processing. Nature 456, 464–469, doi: 10.1038/nature07488 (2008).

10 Hafner, M. et al. Transcriptome-wide identification of RNA-binding protein and microRNA target sites by PAR-CLIP. Cell 141, 129–141, doi: 10.1016/j.cell.2010.03.009 (2010).

11 Van Nostrand, E. L. et al. Robust transcriptome-wide discovery of RNA-binding protein binding sites with enhanced CLIP (eCLIP). Nat Methods 13, 508–514, doi: 10.1038/nmeth.3810 (2016).

12 König, J. et al. iCLIP reveals the function of hnRNP particles in splicing at individual nucleotide resolution. Nat Struct Mol Biol 17, 909–915, doi: 10.1038/nsmb.1838 (2010).

13 Jin, H. et al. TRIBE editing reveals specific mRNA targets of eIF4E-BP in Drosophila and in mammals. Sci Adv 6, eabb8771, doi: 10.1126/sciadv.abb8771 (2020).

14 Rahman, R., Xu, W., Jin, H. & Rosbash, M. Identification of RNA-binding protein targets with HyperTRIBE. Nat Protoc 13, 1829–1849, doi: 10.1038/s41596-018-0020-y (2018).

15 Xu, W., Rahman, R. & Rosbash, M. Mechanistic implications of enhanced editing by a HyperTRIBE RNA-binding protein. RNA 24, 173–182, doi: 10.1261/rna.064691.117 (2018).

16 Singh, A. et al. Antagonistic roles for Ataxin-2 structured and disordered domains in RNP condensation. Elife 10, doi: 10.7554/eLife.60326 (2021).

17 Luo, W. et al. NonA and CPX Link the Circadian Clockwork to Locomotor Activity in Drosophila. Neuron 99, 768–780 e763, doi: 10.1016/j.neuron.2018.07.001 (2018).

18 Herzog, J. J. et al. TDP-43 dysfunction restricts dendritic complexity by inhibiting CREB activation and altering gene expression. Proc Natl Acad Sci U S A 117, 11760–11769, doi: 10.1073/pnas.1917038117 (2020).

19 Liu, M. et al. TRIBE Uncovers the Role of Dis3 in Shaping the Dynamic Transcriptome in Malaria Parasites. Front Cell Dev Biol 7, 264, doi: 10.3389/fcell.2019.00264 (2019).

20 Arribas-Hernandez, L. et al. Principles of mRNA targeting via the Arabidopsis m(6)A-binding protein ECT2. Elife 10, doi: 10.7554/eLife.72375 (2021).

21 Lehrbach, N. J. et al. LIN-28 and the poly(U) polymerase PUP-2 regulate let-7 microRNA processing in Caenorhabditis elegans. Nat Struct Mol Biol 16, 1016–1020, doi: 10.1038/nsmb.1675 (2009).

22 Soleymanjahi, S., Blanc, V. & Davidson, N. APOBEC1 mediated C-to-U RNA editing: target sequence and trans-acting factor contribution to 177 RNA editing events in 119 murine transcripts in-vivo. RNA, doi:10.1261/rna.078678.121 (2021).

23 Abruzzi, K. C., Ratner, C. & Rosbash, M. Comparison of TRIBE and STAMP for identifying targets of RNA binding proteins in human and <em>Drosophila</em> cells. bioRxiv, 2023.2002.2003.527025, doi: 10.1101/2023.02.03.527025 (2023).

24 Rutter, J. & Hughes, A. L. Power(2): the power of yeast genetics applied to the powerhouse of the cell. Trends Endocrinol Metab 26, 59–68, doi: 10.1016/j.tem.2014.12.002 (2015).

25 Mitchell, S. F., Jain, S., She, M. & Parker, R. Global analysis of yeast mRNPs. Nat Struct Mol Biol 20, 127–133, doi: 10.1038/nsmb.2468 (2013).

26 Denisenko, O. & Bomsztyk, K. Epistatic interaction between the K-homology domain protein HEK2 and SIR1 at HMR and telomeres in yeast. J Mol Biol 375, 1178–1187, doi: 10.1016/j.jmb.2007.11.001 (2008).

27 Aronov, S. et al. mRNAs encoding polarity and exocytosis factors are cotransported with the cortical endoplasmic reticulum to the incipient bud in Saccharomyces cerevisiae. Mol Cell Biol 27, 3441–3455, doi: 10.1128/mcb.01643-06 (2007).

28 Hasegawa, Y., Irie, K. & Gerber, A. P. Distinct roles for Khd1p in the localization and expression of bud-localized mRNAs in yeast. Rna 14, 2333–2347, doi: 10.1261/rna.1016508 (2008).

29 Shepard, K. A. et al. Widespread cytoplasmic mRNA transport in yeast: identification of 22 bud-localized transcripts using DNA microarray analysis. Proc Natl Acad Sci U S A 100, 11429–11434, doi: 10.1073/pnas.2033246100 (2003).

30 Wolf, J. J. et al. Feed-forward regulation of a cell fate determinant by an RNA-binding protein generates asymmetry in yeast. Genetics 185, 513–522, doi: 10.1534/genetics.110.113944 (2010).

31 Mauchi, N., Ohtake, Y. & Irie, K. Stability control of MTL1 mRNA by the RNA-binding protein Khd1p in yeast. Cell Struct Funct 35, 95–105, doi: 10.1247/csf.10011 (2010).

32 Das, S., Vera, M., Gandin, V., Singer, R. H. & Tutucci, E. Intracellular mRNA transport and localized translation. Nat Rev Mol Cell Biol 22, 483–504, doi: 10.1038/s41580-021-00356-8 (2021).

33 Paquin, N. et al. Local activation of yeast ASH1 mRNA translation through phosphorylation of Khd1p by the casein kinase Yck1p. Mol Cell 26, 795–809, doi: 10.1016/j.molcel.2007.05.016 (2007).

34 Irie, K. et al. The Khd1 protein, which has three KH RNA-binding motifs, is required for proper localization of ASH1 mRNA in yeast. EMBO J 21, 1158–1167, doi: 10.1093/emboj/21.5.1158 (2002).

35 Singer-Kruger, B. & Jansen, R. P. Here, there, everywhere. mRNA localization in budding yeast. RNA Biol 11, 1031–1039, doi: 10.4161/rna.29945 (2014).

36 Trautwein, M., Dengjel, J., Schirle, M. & Spang, A. Arf1p provides an unexpected link between COPI vesicles and mRNA in Saccharomyces cerevisiae. Mol Biol Cell 15, 5021–5037, doi: 10.1091/mbc.e04-05-0411 (2004).

37 Sezen, B., Seedorf, M. & Schiebel, E. The SESA network links duplication of the yeast centrosome with the protein translation machinery. Genes Dev 23, 1559–1570, doi: 10.1101/gad.524209 (2009).

38 Lang, B. D., Li, A., Black–Brewster, H. D. & Fridovich-Keil, J. L. The brefeldin A resistance protein Bfr1p is a component of polyribosome-associated mRNP complexes in yeast. Nucleic Acids Res 29, 2567–2574, doi: 10.1093/nar/29.12.2567 (2001).

39 Simpson, C. E., Lui, J., Kershaw, C. J., Sims, P. F. & Ashe, M. P. mRNA localization to P-bodies in yeast is bi-phasic with many mRNAs captured in a late Bfr1p-dependent wave. J Cell Sci 127, 1254–1262, doi: 10.1242/jcs.139055 (2014).

40 Kuttan, A. & Bass, B. L. Mechanistic insights into editing-site specificity of ADARs. Proc Natl Acad Sci U S A 109, E3295–3304, doi: 10.1073/pnas.1212548109 (2012).

41 Lewis, H. A. et al. Sequence-specific RNA binding by a Nova KH domain: implications for paraneoplastic disease and the fragile X syndrome. Cell 100, 323–332, doi: 10.1016/s0092-8674(00)80668-6 (2000).

42 Hogan, D. J., Riordan, D. P., Gerber, A. P., Herschlag, D. & Brown, P. O. Diverse RNA-binding proteins interact with functionally related sets of RNAs, suggesting an extensive regulatory system. PLoS Biol 6, e255, doi: 10.1371/journal.pbio.0060255 (2008).

43 Eugster, A., Frigerio, G., Dale, M. & Duden, R. The alpha- and beta’-COP WD40 domains mediate cargo-selective interactions with distinct di-lysine motifs. Mol Biol Cell 15, 1011–1023, doi: 10.1091/mbc.e03-10-0724 (2004).

44 Samanfar, B. et al. A global investigation of gene deletion strains that affect premature stop codon bypass in yeast, Saccharomyces cerevisiae. Mol Biosyst 10, 916–924, doi: 10.1039/c3mb70501c (2014).

45 Gui, Q. et al. Transcriptome Analysis in Yeast Reveals the Externality of Position Effects. Mol Biol Evol 38, 3294–3307, doi: 10.1093/molbev/msab104 (2021).

46 Gibson, D. G. et al. Enzymatic assembly of DNA molecules up to several hundred kilobases. Nat Methods 6, 343–345, doi: 10.1038/nmeth.1318 (2009).

47 Gietz, R. D. & Schiestl, R. H. High-efficiency yeast transformation using the LiAc/SS carrier DNA/PEG method. Nat Protoc 2, 31–34, doi: 10.1038/nprot.2007.13 (2007).

48 Palme, J., Wang, J. & Springer, M. Variation in the modality of a yeast signaling pathway is mediated by a single regulator. Elife 10, doi: 10.7554/eLife.69974 (2021).

49 Green, M. R. & Sambrook, J. Molecular Cloning: A Laboratory Manual (Fourth Edition). (Cold Spring Harbor Laboratory Press, 2012).

50 Alpern, D. et al. BRB-seq: ultra-affordable high-throughput transcriptomics enabled by bulk RNA barcoding and sequencing. Genome Biol 20, 71, doi: 10.1186/s13059-019-1671-x (2019).

51 Bailey, T. L., Johnson, J., Grant, C. E. & Noble, W. S. The MEME Suite. Nucleic Acids Res 43, W39–49, doi: 10.1093/nar/gkv416 (2015).

52 Wu, T. et al. clusterProfiler 4.0: A universal enrichment tool for interpreting omics data. Innovation (Camb) 2, 100141, doi: 10.1016/j.xinn.2021.100141 (2021).

53 Jan, C. H., Williams, C. C. & Weissman, J. S. Principles of ER cotranslational translocation revealed by proximity-specific ribosome profiling. Science 346, 1257521, doi: 10.1126/science.1257521 (2014).

